# Reduced CSF1R expression in myeloid cells has limited impact on chronic lymphocytic leukemia progression

**DOI:** 10.1101/2025.10.07.680920

**Authors:** Natascha Rosen, Guillermo Rodriguez-Real, Viktoria Kohlhas, Thanh-Tung Truong, Alexander F. vom Stein, Maximilian Koch, Sebastian Reinartz, Sumiya Iqbal, Shaista Ilyas, Sanjay Mathur, Nina Reinart, Phuong-Hien Nguyen, Michael Hallek

## Abstract

Targeting the colony-stimulating factor 1 receptor (CSF1R) to remove tumor-associated macrophages is being explored as cancer therapy. This strategy may be relevant for chronic lymphocytic leukemia (CLL), which strongly depends on support from myeloid cells. However, it is unclear how CSF1R expression affects the CLL microenvironment and leukemic progression. To examine this question, we created CLL mice with *Csf1r* haploinsufficiency to investigate changes in myeloid cells, Csf1r expression, and leukemia progression. Reducing Csf1r expression on circulating monocytes did not change the overall numbers of monocytes or macrophages in blood and lymphoid tissues. Mice with lower Csf1r levels had less leukemia during early disease, but this effect faded with disease progression, and their overall survival was similar to controls. Furthermore, reduced Csf1r expression on macrophages did not affect the survival or migration of patient-derived CLL cells.

In summary, our results show that while the CSF1R pathway is important for maintaining the myeloid cells that support CLL, simply reducing CSF1R expression has only a limited effect on disease progression. Attempts to target the CSF1R for leukemic therapy might benefit from a stronger depletion of macrophages or the combination with other agents.

## Introduction

Chronic lymphocytic leukemia (CLL) is the most common leukemia in the Western world. It is characterized by the monoclonal expansion and accumulation of mature, yet immunodeficient, CD5^+^ B cells in the blood, lymphoid tissues and bone marrow (Hallek, 2025).

Although CLL cells persist long-term in patients, they rapidly undergo apoptosis in vitro (Collins et al., 1989). This is explained by the importance of the tumor microenvironment (TME), which provides essential survival signals (Vom Stein, Hallek, & Nguyen, 2024). Among TME components, macrophages play a central role in CLL progression and treatment responses (Jestrabek, Kohlhas, Hallek, & Nguyen, 2024). Increased monocyte counts in blood correlate with poor prognosis (Hanna, Ozturk, & Seiffert, 2019; Ten Hacken & Burger, 2016). Peripheral monocytes, upon contact with leukemic cells, differentiate into nurse-like cells (NLCs) that prevent apoptosis and support CLL cell viability (Burger et al., 2000). These supportive features can be induced by leukemia-derived factors such as HMGB1 and NAMPT (Audrito et al., 2015; Jia et al., 2014). In return, leukemia-associated macrophages contribute to CLL cell survival and disease progression via secretion of soluble factors such as CD14 and MIF, inducing kinase signaling such as JAK/STAT and MAPK, and enhancing leukemic NF-kB signaling (Hanna et al., 2019; Viktoria Kohlhas et al., 2025; Nguyen et al., 2016; Nguyen, Niesen, & Hallek, 2019; Reinart et al., 2013; Seiffert et al., 2010), making them attractive therapeutic targets.

A central regulator of TME organization is colony-stimulating factor 1 (CSF-1), a cytokine elevated in CLL patients (Janowska-Wieczorek et al., 1991). CSF-1 is critical for the survival, proliferation, and differentiation of mononuclear phagocytes (Stanley, Guilbert, Tushinski, & Bartelmez, 1983), acting exclusively through the CSF1 receptor (CSF1R) (Dai et al., 2002). CSF1R is expressed on monocytes, macrophages, and several other cell types, including osteoclasts, neurons, and cells of the female reproductive tract (Violeta Chitu et al., 2015). In mice, complete Csf1r deficiency leads to substantial loss of macrophages, skeletal abnormalities including toothlessness, growth retardation, and reduced survival, with strain-dependent variability (V. Chitu, Gokhan, Nandi, Mehler, & Stanley, 2016; Dai et al., 2002).

In normal B-cell development, CSF1R is only expressed in early precursors (Zriwil et al., 2016) and is repressed during maturation (Tagoh et al., 2006). In CLL, CSF1R is not aberrantly expressed on leukemia cells (Edwards et al., 2018), and its levels appear similar in monocytes and NLCs (Polk et al., 2016). However, preclinical studies have shown that blocking CSF1R can disrupt macrophage support and trigger apoptosis in CLL cells. For example, treatment with anti-Csf1r antibody in Rag2^*-/-*^ γc^*-/-*^ mice transplanted with MEC-1 cells reduced macrophage numbers, decreased tumor burden, and caused delocalization of CLL cells from tissues to blood, enhancing their sensitivity to anti-CD20 therapy (Galletti et al., 2016). Similar effects were observed in vitro using small-molecule inhibitors (GW-2580, ARRY-382) or antibodies targeting CSF1R (Galletti et al., 2016; Polk et al., 2016). These studies primarily aimed to deplete macrophages, but did not investigate whether modulating CSF1R expression levels could influence disease development.

To address this gap, we investigated the role of CSF1R expression level in CLL by adopting a genetic approach: we used heterozygous *Csf1r*^*+/-*^ mice, which are viable despite reduced Csf1r expression, thus avoiding the embryonic lethality observed in Csf1r^*-/-*^ animals (Dai et al., 2002). *Csf1r*^*+/-*^ mice were crossed with the *Eµ-TCL1* transgenic mice, serving as a model for CLL (Autio et al., 2020; Koch et al., 2020), to examine the effects of different Csf1r expression levels in a physiologically relevant setting.

## Material and methods

### CLL samples

Primary CLL samples were collected from peripheral blood with informed consent, following the Declaration of Helsinki and approved by the Institutional Ethics Committee (no. 11-319) of the University Hospital Cologne. PBMCs and CLL cells were cryopreserved at −150 °C and freshly thawed for each assay.

### Mouse experiments

*Csf1r*^*+/-*^ mice (Dai et al., 2002) on C57BL6J x C3Heb/FeJ background (kind gift from E. Richard Stanley, NYC, USA) were crossed with *Eµ-TCL1* transgenic mice (Bichi et al., 2002) on C57BL/6J background. The resulting *TCL1*^*tg/wt*^ *Csf1r*^*+/-*^ and control *TCL1*^*tg/wt*^ *Csf1r*^*+/+*^ mice were housed in a conventional pathogen-free facility at the University Hospital Cologne.

For survival experiments, mice were monitored from birth until death or euthanasia according to approved animal welfare guidelines. Blood was collected at months 2, 4, 6, 8, 10, and 12 via tail vein incision into EDTA-coated microvettes (Sarstedt) and kept refrigerated until analysis (V. Kohlhas, Hallek, & Nguyen, 2020; Nguyen et al., 2016; Reinart et al., 2013).

All mouse procedures were approved by the North Rhine-Westphalia state authorities (LANUV: 84-02.04.2014.A146; 81-02.04.2019.A009).

### Hematologic Analysis

Complete blood counts were obtained using a Sysmex XE-5000 hematology analyzer (Sysmex Corporation), which quantifies white blood cells (WBCs) via impedance and optical scatter without antibody labeling. For flow cytometry, red blood cells (RBCs) were lysed with ACK buffer, and the remaining leukocytes were washed and resuspended in PBS.

### Flow cytometry

Single-cell suspensions were stained with fluorochrome-conjugated antibodies (Supplementary Table 1) as previously described (V. Kohlhas et al., 2020; Nguyen et al., 2016; Reinart et al., 2013). Data were acquired using a MACSQuant VYB cytometer (Miltenyi Biotec) and analyzed with Kaluza (Beckman Coulter) or FlowJo (BD Pharmingen). Mean fluorescence intensity (MFI) was calculated as indicated in the respective figure legends.

### Immunohistochemistry

Spleen tissues from 8- to 10-month-old mice were fixed in 4% buffered paraformaldehyde and paraffin-embedded. Sections were stained with antibodies listed in Supplementary Table 2, following the manufacturers’ instructions. Signal detection was performed using the Histofine® Simple Stain Mouse MAX PO polymer-based system (Nichirei Biosciences).

### Cell proliferation assay

Peripheral blood, bone marrow and spleen were collected from 8- to 10-month-old mice. To prepare single cell suspensions from bone marrow, both tibia and femur were flushed with PBS. Spleen cells were passed through a 100-μm cell strainer (Greiner Bio-One) and RBCs were lysed using ACK lysing buffer, washed and resuspended in PBS. Intracellular Ki-67 staining was performed using IntraPrep Permeabilization Reagent (Beckman Coulter) following manufacturer’s instructions. For specific antibodies, see Supplementary Table 1.

### Cell culture

MacCsf1r^+/+^ and MacCsf1r^-/-^ murine macrophage cell lines (Yu et al., 2008) were kindly provided by E. Richard Stanley (NYC, USA) and cultured in DMEM (Gibco) containing 10% FBS (PAN-Biotech), 1% Pen Strep (Gibco) and mouse GM-CSF (Biolegend) at concentrations indicated for each experiment below.

THP-1 monocyte cell line (RRID:CVCL_0006) was purchased from DSMZ (ACC16) and cultured in RPMI (Gibco) supplemented with 10% FBS and 1% Pen Strep.

### Generation of *CSF1R* knockout THP-1 cells via CRISPR/Cas9

The THP-1 *CSF1R*^*-/-*^ cells were generated in two steps. First, THP-1 cells (1 × 10^6^) were infected twice with third-generation lentiviral vectors encoding Cas9-eGFP (Thermo Fisher) at MOI 5 by spin infection (800 rpm, 120 min, 37 °C), and single-cell clones with high eGFP expression were isolated by FACS.

For *CSF1R* knockout, cells were transduced with lentiviral particles containing four gRNAs targeting human *CSF1R* (Thermo Fisher; gRNA-1: CAACGCTACCTTCCAAAACA; gRNA-2: GAACGTGCTAGCACAGGAGG; gRNA-3: AACGGTGACCTTGCGATGTG; gRNA-4: GGACACTGGGCTCTATCACT). Scramble controls were generated using a non-targeting gRNA (scr-gRNA: GTACGTCGGTATAACTCCTC; Invitrogen LentiArray™).

Cas9-expressing THP-1 cells (1 × 10^4^ per well) were seeded into 96-well plates and transduced at MOI 10 using spin infection (800 rpm, 90 min, 37 °C). After 72 hours, cells were washed and selected in RPMI with 0.75 μg/mL puromycin (Carl Roth) for over 7 days, replenishing medium after 3 days.

### Western blot

Cells were lysed in RIPA buffer with 1 mM PMSF (Cell Signaling), incubated on ice for 1 hour, centrifuged, and supernatants stored at −80 °C. Immunoblots were performed per manufacturer’s instructions using NuPAGE 4–12% Bis-Tris gels (Thermo Scientific) and semi-dry transfer to nitrocellulose membranes (GE Healthcare). Signals were detected by immunofluorescence on a LiCor Odyssey CXl. Antibodies are listed in Supplementary Table 3.

### Synthesis of Doxorubicin-loaded liposomes (Lipo-Doxo)

Doxorubicin-loaded liposomes were prepared using the thin-film hydration method with DPPC, cholesterol, and DSPE-PEG(2000)-NH_2_ dissolved in chloroform. After solvent evaporation under reduced pressure and vacuum drying (2 hours), the lipid film was hydrated with 250 mM ammonium hydrogen phosphate ((NH_4_)_2_HPO_4_) buffer at 60 °C, followed by five freeze–thaw cycles and extrusion through 200 nm polycarbonate membranes to produce unilamellar vesicles.

The external buffer was exchanged for PBS (pH 7.4) via Sephadex G-50 size exclusion chromatography (SEC) to establish a pH gradient. Doxorubicin hydrochloride was added at a 1:10 drug-to-lipid molar ratio and incubated at 60 °C for 1 hour for active loading. The unencapsulated drug was removed by a second SEC step. The final formulation had a phospholipid concentration of 570.70 µg/mL (Stewart assay) and a doxorubicin loading capacity of 37.19% (HPLC). The two SEC steps ensured purity but contributed to partial lipid dilution.

### CLL co-culture and viability assays

For MacCsf1r co-cultures, 1 × 10^5^ cells were seeded in 24-well plates and incubated overnight before adding 5 × 10^5^ cryopreserved CLL cells. Culture medium was supplemented with 200 ng/mL GM-CSF.

In THP-1 co-cultures, 1 × 10^5^ THP-1 cells were differentiated with 100 ng/mL phorbol 12-myristate 13-acetate (PMA; Sigma-Aldrich) for 48 h, washed to remove PMA, and co-cultured with 5 × 10^5^ patient-derived CLL cells.

For Lipo-Doxo toxicity assays, 3 × 10^5^ PBMCs from three healthy donors were seeded in 96-well plates and treated with 0.1 μM or 1 μM Lipo-Doxo. After 24 h, viability of total cells and immune subsets was analyzed by flow cytometry.

In mixed PBMC–CLL cultures, 5 × 10^6^ PBMCs (±1 μM Lipo-Doxo, 24 h) were co-cultured with 5 × 10^5^ CLL cells from two patients in 24-well plates. PBMCs were washed and resuspended in RPMI before being added to CLL cultures. Monocultures of CLL cells served as controls. At 24, 48 and 72 h, 200μl of cell suspension from each well was transferred to 96 well-plates for viability analysis. All conditions were in technical triplicates.

CLL viability was measured by flow cytometry using Annexin V antibody (BioLegend) on a MACSQuant X cytometer (Miltenyi Biotec), analyzed with FlowJo (BD Pharmingen).

### CLL migration assays towards macrophages

MacCsf1r cells (1 × 10^5^) were seeded in 24-well plates. After 24 h, medium was replaced with RPMI + 10% FBS, 1% Pen/Strep, and 110 ng/mL GM-CSF. After 20 h, serum-starved CLL cells (1 × 10^6^, 2 h) were added to transwell inserts (5 μm pores, Corning). After 4 h, migrated cells were counted by flow cytometry (MACSQuant X, Miltenyi Biotec) and analyzed with FlowJo (BD Pharmingen).

For THP-1 cells, 1 × 10^5^ cells were differentiated into macrophages by stimulation with 100 ng/mL PMA for 48 hours. After differentiation, THP-1 macrophages were washed twice with PBS and 1 × 10^6^ CLL cells were added, and the same protocol previously described for MacCsf1r cells was followed.

### Statistical analysis and data presentation

Statistical analyses were performed using GraphPad Prism 8.4.0. Unless stated otherwise, the Mann–Whitney test was used. Dot plots show medians; line graphs show mean ± SEM. P-values are indicated as: * ≤ 0.05, ** ≤ 0.01, *** ≤ 0.001, **** ≤ 0.0001. Figures were prepared using GraphPad Prism and Adobe Illustrator.

## Results

### Monocyte depletion impairs support of CLL cell survival

To assess whether circulating monocytes support CLL survival, we conducted co-cultured of patient-derived CLL cells with peripheral blood mononuclear cells (PBMCs) from healthy donors over 24, 48 and 72 hours. To deplete monocytes, PBMCs were treated with liposomes encapsulating the chemotherapy drug doxorubicin (Lipo-Doxo), which significantly reduced the viability of the monocyte subset (CD14+CD3^−^CD4^−^CD8^−^CD20^−^) compared to the total PBMC population at 1 µM (Supplementary Figure 1A). In co-culture with CLL cells, PBMCs pre-treated with 1 µM Lipo-Doxo exhibited a significantly reduced capacity to support CLL cell survival compared to the DMSO-treated controls (Supplementary Figure 1B). Having confirmed the critical role of monocytes in supporting CLL cell survival, we next investigated the role of CSF1R, primarily expressed on monocytes and macrophages, in this crosstalk.

### Heterozygous deletion of *Csf1r* significantly reduces Csf1r expression on peripheral myeloid cells

To modulate *Csf1r* expression independently of myeloid cell depletion, we examined the myeloid compartment of *Csf1r*^*+/-*^ mice, which show reduced Csf1r levels (Dai et al., 2002). Longitudinal blood analysis revealed comparable counts of CD11b+blood monocytes and total white blood cells (WBCs) between *Csf1r*^*+/-*^ and wild-type mice (Figure 1A, 1B). Likewise, CD11b+myeloid cell frequencies in spleen and bone marrow remained unchanged (Figure 1C).

**Figure 1.**
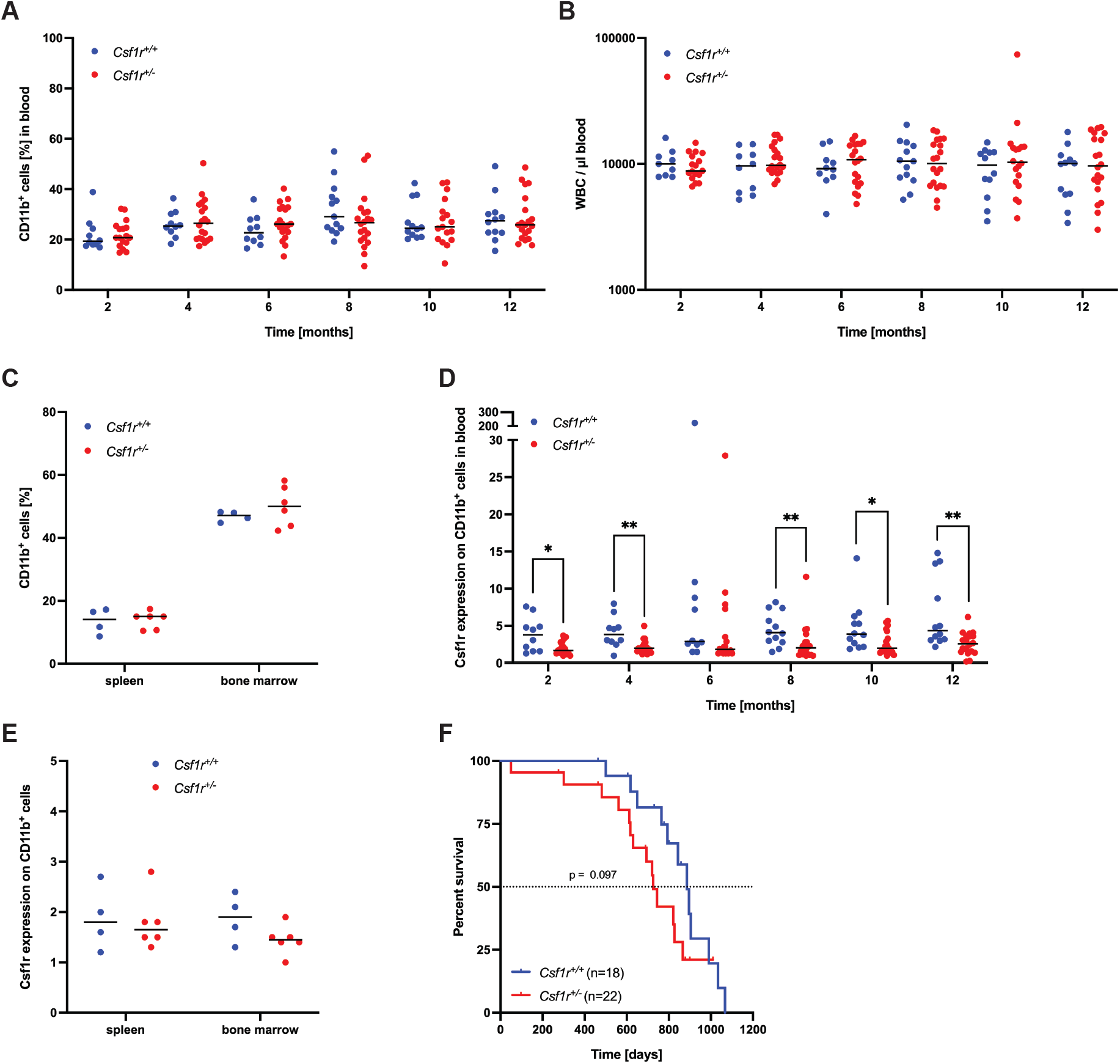
Heterozygous deletion of *Csf1r* significantly reduced Csf1r expression on peripheral myeloid cells. A) Longitudinal analysis over one year showing monocyte populations (CD11b+ cells) in peripheral blood from *Csf1r*^*+/+*^ and *Csf1r*^*+/-*^ mice. B) White blood cell (WBC) counts for *Csf1r*^*+/+*^ and *Csf1r*^*+/-*^ mice, measured using the XE-5000 hematology analyzer. C) Flow cytometry analysis of CD11b^+^ cells in the spleen and bone marrow of 8–10-month-old *Csf1r*^*+/+*^ and *Csf1r*^*+/-*^ mice. D) Csf1r expression on CD45^+^ CD11b^+^ peripheral blood cells over one year, shown as the mean fluorescence intensity (MFI) ratio normalized to the isotype control. E) Protein expression of Csf1r on CD11b^+^ cells from the spleen and bone marrow of 8–10-month-old mice, expressed as the MFI ratio normalized to the FMO (fluorescence minus one). F) Kaplan–Meier survival curves for *Csf1r*^*+/+*^ and *Csf1r*^*+/-*^ mice. Statistical comparison was performed using the log-rank (Mantel–Cox) test.

Despite normal cell counts, *Csf1r*^*+/-*^ mice displayed significantly reduced Csf1r expression on circulating CD11b+monocytes (Figure 1D), while expression in spleen and bone marrow myeloid cells showed no significant differences (Figure 1E). Finally, *Csf1r*^*+/-*^ mice showed a slight but non-significant reduction in lifespan, living an average of 160 days shorter than their wild-type littermates (Figure 1F).

To assess the impact of reduced monocytic Csf1r expression on CLL development, we crossed *Csf1r*^*+/-*^ with *Eµ-TCL1* transgenic mice to generate *TCL1*^*tg/wt*^ *Csf1r*^*+/+*^ and *TCL1*^*tg/wt*^ *Csf1r*^*+/-*^ cohorts. Consistent with *Csf1r*^*+/-*^ mice, longitudinal blood analysis revealed no significant difference in monocyte counts in *TCL1*^*tg/wt*^ mice regardless of their *Csf1r* genotype (Figure 2A). Similarly, myeloid cell frequencies in the spleen and bone marrow remained stable (Figures 2B, 2C). As observed in *Csf1r*^*+/-*^ mice, Csf1r expression on blood monocytes was significantly reduced in *TCL1*^*tg/wt*^ *Csf1r*^*+/-*^ mice (Figure 2D), with no change in lymphoid tissue myeloid cells (Figure 2E). As expected, CLL cells (Edwards et al., 2018) and healthy B cells (Zriwil et al., 2016) did not express Csf1r (Figure 2F).

**Figure 2.**
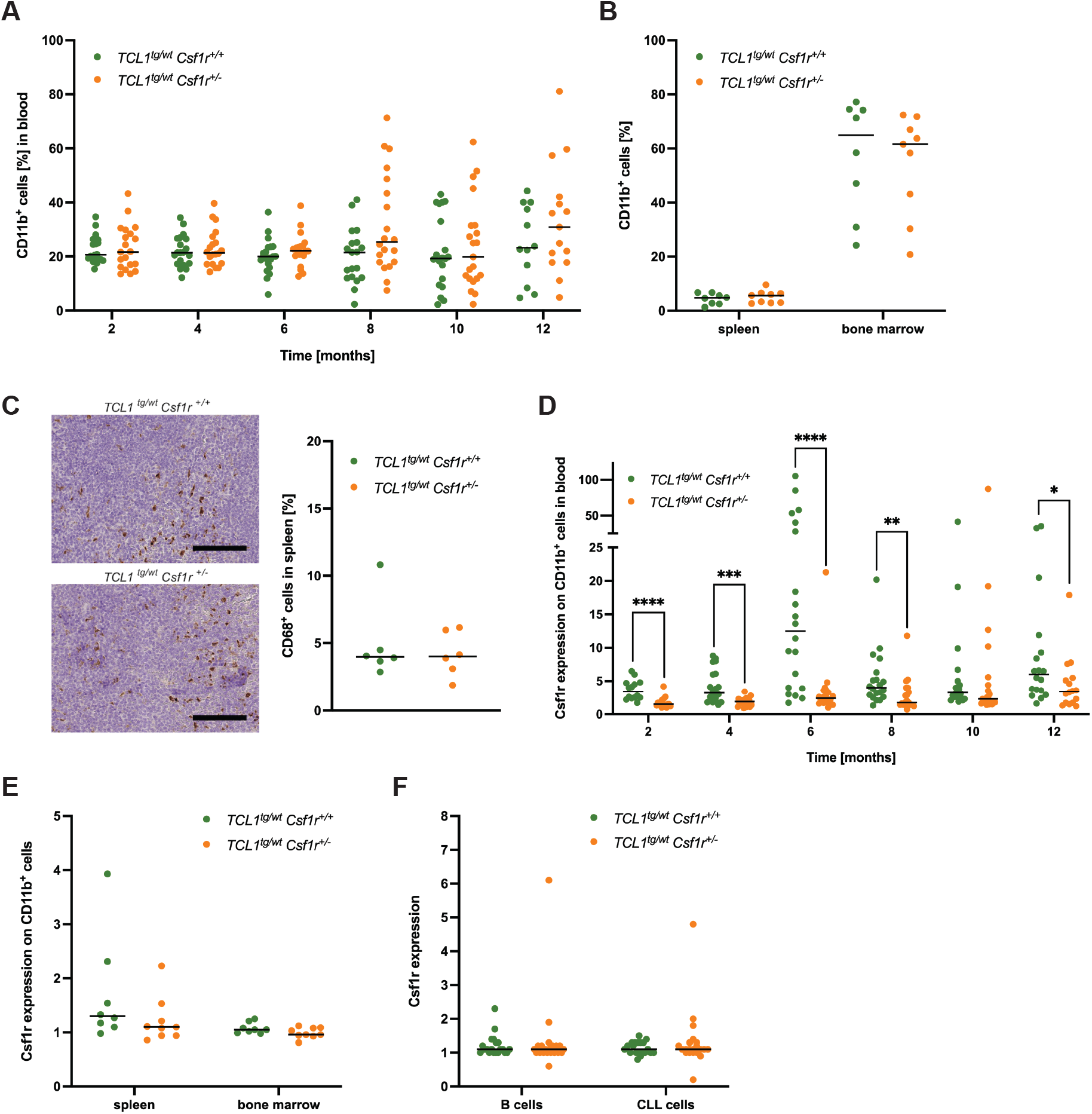
Heterozygous deletion of *Csf1r* in the *TCL1*^*tg/wt*^ mice reiterated the reduced Csf1r expression on peripheral myeloid cells. A) Longitudinal peripheral blood analysis over one year showing CD11b+ monocyte populations in *TCL1*^*tg/wt*^ *Csf1r*^*+/+*^ and *TCL1*^*tg/wt*^ *Csf1r*^*+/-*^ mice. B) Flow cytometry analysis of CD11b^+^ cells in the spleen and bone marrow of 8–10-month-old *TCL1*^*tg/wt*^ *Csf1r*^*+/+*^ and *TCL1*^*tg/wt*^ *Csf1r*^*+/-*^ mice. C) Representative immunohistochemical staining of CD68^+^ cells in spleen sections (scale bars = 100 µm) and quantification for individual mice. D) Csf1r expression on CD11b^+^ cells in blood over one year, expressed as the MFI ratio normalized to the isotype control.. E) Csf1r expression on CD11b^+^ cells in the spleen and bone marrow, in 8-10-month-old mice, expressed as the MFI ratio normalized to the FMO. F) Flow cytometry analysis of Csf1r expression on CD45^+^ CD19^+^ CD5^-^ B cells and CD45^+^ CD19^+^ CD5^+^ CLL cells in peripheral blood, from 8-10-month-old mice; MFI values were calculated as in (D).

### Loss of one *Csf1r* allele reduced leukemic burden in the peripheral blood of *TCL1*^*tg/wt*^ mice without affecting tissue infiltration

Longitudinal blood analysis over 12 months showed significantly reduced CLL burden at month 8 in *TCL1*^*tg/wt*^ *Csf1r*^*+/-*^ mice (Mean ± SEM = 31.07 ± 4.806% CLL cells in blood) compared to *TCL1*^*tg/wt*^ *Csf1r*^*+/+*^ mice (Mean ± SEM = 45.74 ± 4.566% CLL cells in blood). This difference persisted, albeit less markedly, at 12 months of age (*TCL1*^*tg/wt*^ *Csf1r*^*+*^*/*^*+*^: 42.20 ± 7.376%; *TCL1*^*tg/wt*^ *Csf1r*^*+*^*/*^*−*^: 35.41 ± 6.679%) (Figure 3A). Although the absolute number of leukemic cells in *TCL1*^*tg/wt*^ *Csf1r*^*+/-*^ mice followed the same trend, the difference did not reach statistical significance (Figure 3B), and WBC counts were comparable between genotypes (Figure 3C). Furthermore, Ki-67 staining showed no differences in CLL proliferation across blood, spleen, and bone marrow samples between *TCL1*^*tg/wt*^ *Csf1r*^*+/+*^ and *TCL1*^*tg/wt*^ *Csf1r*^*+/-*^ mice (Figure 3D), suggesting minimal influence of Csf1r expression on CLL proliferation.

**Figure 3.**
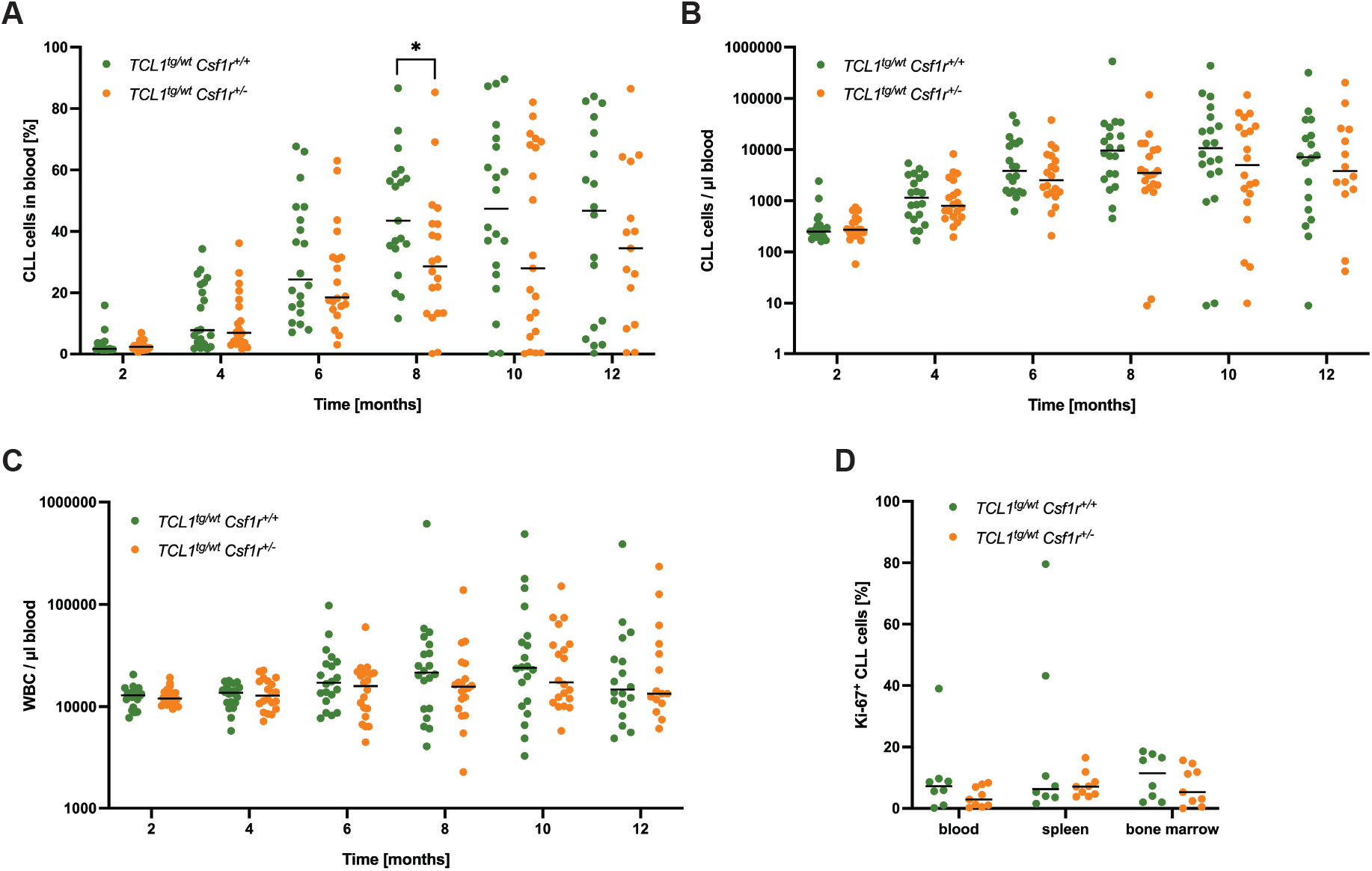
Loss of one *Csf1r* allele reduced leukemic burden in blood but not in lymphoid tissues of *TCL1*^*tg/wt*^ mice. A) Longitudinal flow cytometric analysis of CD45^+^ CD19^+^ CD5^+^ CLL cells in the peripheral blood of *TCL1*^*tg/wt*^ *Csf1r*^*+/+*^ and *TCL1*^*tg/wt*^ *Csf1r*^*+/-*^ mice over one year. B) Absolute numbers of CLL cells, calculated from the WBC counts. C) WBC counts measured with the XE-5000 hematology analyzer. D) Proliferating CLL cells (CD19^+^ CD5^+^ Ki-67^+^) in blood, spleen and bone marrow from 8–10-month-old mice.

Further analysis of lymphoid tissue infiltration between months 8 and 10 – the time point of significant reduction in peripheral blood CLL cells in *TCL1*^*tg/wt*^ *Csf1r*^*+/-*^ mice (Figure 3A) – showed no differences between genotypes. Spleen and liver weights were comparable between genotypes (Figure 4A), and leukemic infiltration in spleen and bone marrow was similar (Figure 4B). Immunohistochemistry staining again confirmed comparable spleen infiltration (Figure 4C). Moreover, overall survival was also identical, with both groups living an average of 358 days (Figure 4D). Taken together, these results indicate that *Csf1r* haploinsufficiency mainly reduced CLL abundance in peripheral blood of *TCL1*^*tg/wt*^ mice around the disease acceleration phase, without affecting tissue burden, and this effect diminished as leukemia progressed.

**Figure 4.**
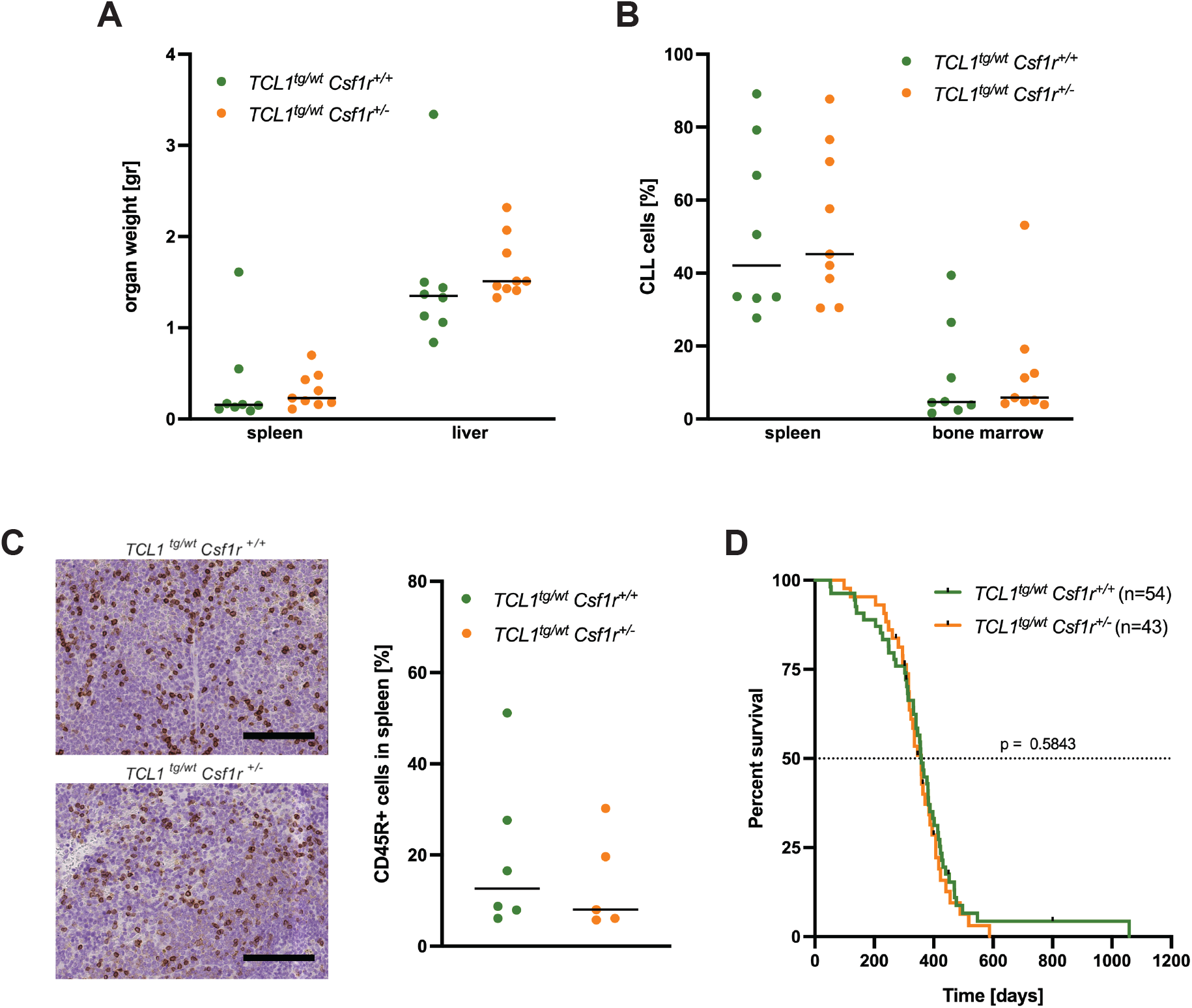
CLL development in lymphoid tissue was not affected by heterozygous *Csf1r* deletion. A) Spleen and liver weights in 8-10-month-old *TCL1*^*tg/wt*^ *Csf1r*^*+/+*^ and *TCL1*^*tg/wt*^ *Csf1r*^*+/-*^ mice. B) CD19^+^ CD5^+^ CLL cell infiltration in the spleen and bone marrow in 8-10-month-old mice. C) Representative immunohistochemical staining of CD45R^+^ cells in spleen sections (scale bars = 100 µm) with quantification for individual 8-10-month-old mice. D) Kaplan–Meier survival curves for *TCL1*^*tg/wt*^ *Csf1r*^*+/+*^ and *TCL1*^*tg/wt*^ *Csf1r*^*+/-*^ mice. Statistical comparison was performed using the log-rank (Mantel–Cox) test.

### CSF1R-deficient macrophages exhibit no significant alteration in their capacity to recruit and support CLL cells in vitro

To further explore the role of Csf1r in macrophage mediated CLL support, we co-cultured patient-derived CLL cells with MacCsf1r macrophages, an immortalized murine cell line lacking Csf1r expression (Yu et al., 2008). As we recently reported (Viktoria Kohlhas et al., 2025), CLL cells cultured without feeder macrophages rapidly underwent apoptosis, whereas both MacCsf1r^+/+^ and MacCsf1r^-/-^ macrophages equally supported CLL cells (Figure 5A).

**Figure 5.**
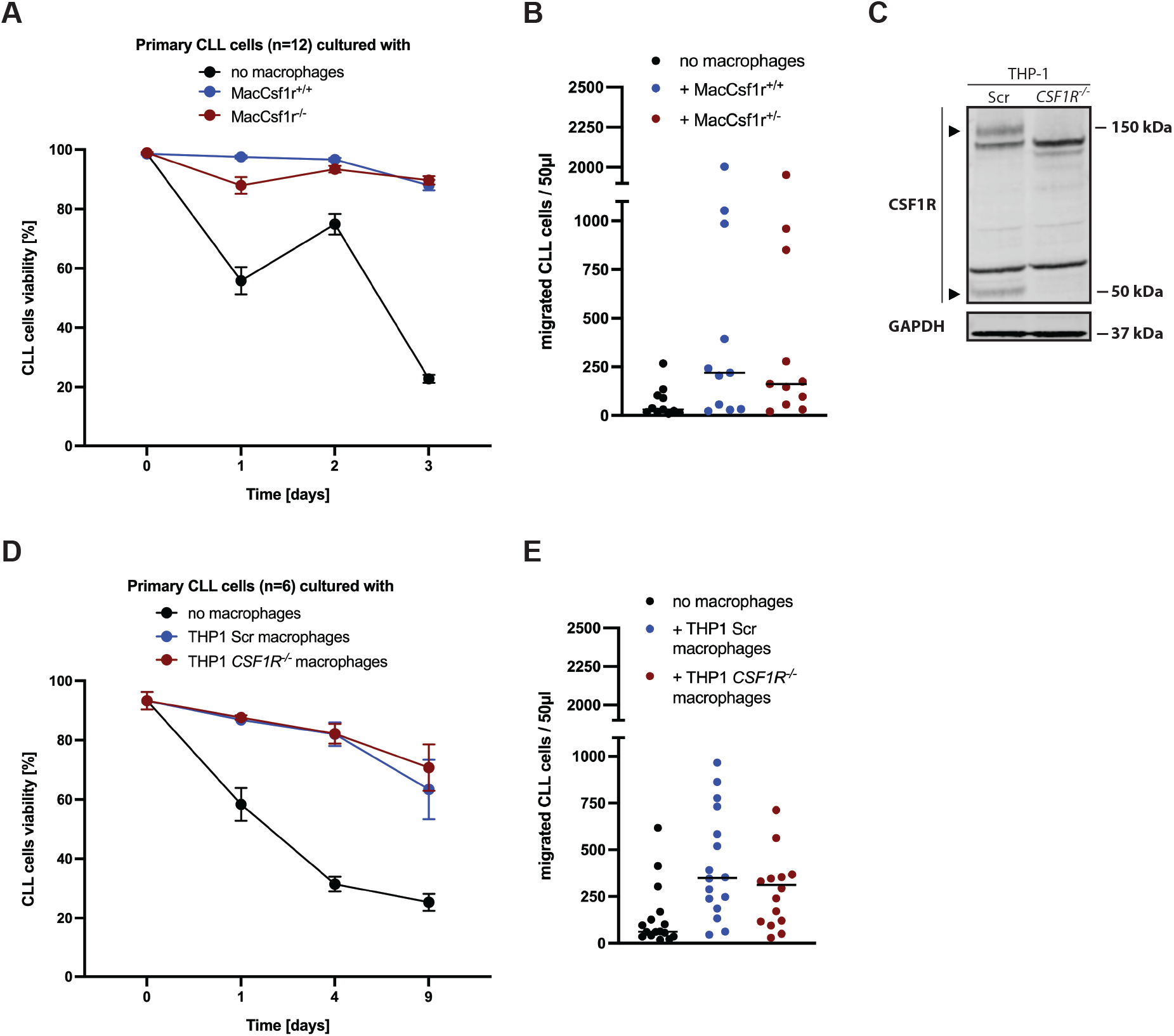
CSF1R-deficient macrophages failed to recruit CLL cells *in vitro*. A) Twelve primary CLL patient samples were co-cultured with either wild-type (MacCsf1r^+/+^), Csf1r-deficient (MacCsf1r^+/-^) macrophages, or without macrophages. CLL cell survival was assessed by Annexin V and 7-AAD staining. Statistical analysis was performed using the Friedman test followed by Dunn’s multiple comparisons test. B) Migration of primary CLL patient cells towards MacCsf1r^+/+^, MacCsf1r^+/-^ macrophages or no macrophages, measured by flow cytometry. Data points represent the mean of technical triplicates; CLL cells were identified by FSC/SSC profiles. C) Validation of *CSF1R* knockout in THP-1 *CSF1R*^*-/-*^ cells by immunoblot analysis of CSF1R protein expression. The ∼150 kDa and ∼50 kDa bands observed in wild-type cells correspond to the mature glycosylated receptor and the cytoplasmic fragment of CSF1R, respectively, and are absent in knockout cells. Remaining bands are considered non-specific. D) Six primary CLL patient samples were co-cultured with control THP-1 macrophages (THP-1 Scr), *CSF1R* knockout THP-1 macrophages (THP-1 *CSF1R*^*-/-*^), or without macrophages. CLL cell survival was measured as described in (A). Statistical analysis was performed using the Kruskal–Wallis test followed by Dunn’s multiple comparisons test. E) Migration of primary CLL cells towards control THP-1 Scr macrophages, THP-1 *CSF1R*^*-/-*^ macrophages or no macrophages, assessed as described in (B).

Similarly, CLL migration toward MacCsf1r macrophages was unaffected by Csf1r depletion (Figure 5B). These results could be confirmed in a fully human co-culture system of CLL cells and THP-1 feeder macrophages (Viktoria Kohlhas et al., 2025; Nguyen et al., 2016). We generated CSF1R deficient THP-1 macrophages, using a CRISPR-Cas9 mediated knockout system (Figure 5C). Upon co-culture with primary human CLL, we observed neither impairment in the ability of CSF1R-knockout macrophages to support CLL survival (Figure 5D), nor in CLL cell recruitment in vitro (Figure 5E).

Together, these in vitro results indicate that CSF1R loss does not impair macrophage-mediated CLL support or recruitment, which is consistent with the unchanged CLL burden in mouse lymphoid tissues (Figures 4B–C), highlighting that the efficacy of CSF1R blockade relies mainly on myeloid cell depletion rather than direct effects of CSF1R expression on leukemia support.

## Discussion

In this study, we show that reducing CSF1R expression on myeloid cells has limited impact on CLL progression both in vivo and in vitro. Using *Csf1r*^*+/-*^ mice, which exhibit reduced receptor expression on blood monocytes while retaining normal myeloid cell counts, we observed a transient reduction in peripheral leukemia burden at around 8 months of age in the *TCL1*^*tg/wt*^ *Csf1r*^*+/-*^ model. However, this effect diminished over time and did not translate into improved survival. This suggests that *Csf1r* haploinsufficiency could only partially disrupt myeloid-mediated support, exerting minimal influence on overall CLL progression. In line with this, the complete loss of CSF1R in macrophages did not significantly affect survival or migration of patient-derived CLL cells in vitro. Together, these results indicate that reducing CSF1R expression level could not overcome the reprogramming of tissue macrophages to support leukemia cells in a CLL mouse model, and therapeutic efficacy of CSF1R-targeted strategies in CLL may depend more on a depletion of myeloid cells than of reduced CSF1R expression levels.

Targeting the CSF1R pathway represents a clinically advanced strategy for modulating the myeloid compartment, with several inhibitors undergoing clinical trials across various malignancies (Gelderblom, Bhadri, et al., 2024; Gelderblom, Razak, et al., 2024; Lei et al., 2020; Spierenburg et al., 2022; Wen, Wang, Guo, & Liu, 2023). The primary goal of these therapies is the depletion of myeloid cells, particularly tumor-associated macrophages (TAMs), whereas the biological effects of solely reducing CSF1R expression remain poorly defined. Our results with liposomal doxorubicin-mediated monocyte depletion showed significantly impaired PBMC-driven CLL cell viability in vitro, confirming the critical role of myeloid support in this disease. Similarly, previous murine studies demonstrated that anti-CSF1R antibodies effectively depleted CLL-associated macrophages, leading to redistribution of leukemic cells into peripheral blood and enhancing sensitivity to therapies (Galletti et al., 2016).

CSF1R inhibitor activity is notably context dependent. For instance, CSF1R blockade failed to effectively deplete TAMs in glioblastoma due to compensatory GM-CSF and IFN-γ signaling from glioma cells (Pyonteck et al., 2013). Additionally, CSF1R inhibition selectively eliminated tissue-resident macrophages but only partially depleted splenic or lung macrophages, highlighting tissue-specific reliance on CSF1R signaling (Bosch et al., 2023; Mohamed et al., 2024). Furthermore, most small-molecule CSF1R inhibitors target other kinases such as FLT, KIT, and VEGFR due to structural homology, complicating attribution of therapeutic effects solely to CSF1R inhibition. Thus, our genetic approaches, like the heterozygous or knockout models used here, provide greater specificity for dissecting CSF1R’s functional role in cancer.

The CSF-1/CSF1R axis is essential for monocyte/macrophage homeostasis, regulating survival, differentiation, and proliferation within the monocytic lineage (Stanley & Chitu, 2014). In CLL, elevated CSF-1 levels likely promote monocyte expansion and polarization toward tumor-supportive macrophages (Polk et al., 2016), correlating with poor prognosis (Gustafson et al., 2012). Although receptor-mediated clearance of CSF-1 complicates detection (Bartocci et al., 1987), *Csf1r* haploinsufficient mice do not exhibit elevated CSF-1 levels (Dai et al., 2002), suggesting unsaturated clearance capacity. Despite reduced receptor levels in blood monocytes, we did not observe significant alterations in myeloid cell populations or CLL proliferation in lymphoid tissues, indicating compensatory mechanisms likely maintain microenvironmental homeostasis.

Interestingly, while confirming that *Csf1r* haploinsufficiency is not dosage compensated (Hume et al., 2020), we observed no significant differences in Csf1r expression between wild-type and *Csf1r*^*+/-*^ myeloid cells in neither spleen nor bone marrow. Likewise, monocyte/macrophage populations and leukemic cell distribution remained stable in lymphoid niches. Notably, the transient reduction in peripheral tumor load at 8 months suggests that during accelerated disease phase CLL cells are more dependent on microenvironmental support, including Csf1r-expressing myeloid cells. As the disease progresses and CLL cells become more autonomous, this dependency appears to diminish, potentially explaining the reduced effect at later stages. This interpretation may also be influenced by compartment-specific Csf1r expression, as reported in earlier studies (Byrne, Guilbert, & Stanley, 1981; Rojo et al., 2019).

Future studies employing cell-type-specific *Csf1r* knockout models (Schreiber et al., 2013; Theurich et al., 2017) could further clarify distinct roles of CSF1R signaling in blood monocytes versus tissue-resident macrophages. Additionally, exploring interactions between CSF1R and other critical pathways such as PI3K/AKT or NF-κB (Guo et al., 2024; Rascio et al., 2021) may elucidate mechanisms of resistance and suggest novel combinatorial approaches.

Clinically, small-molecule CSF1R inhibitors such as pexidartinib and vimseltinib are effective and FDA-approved in tenosynovial giant cell tumor (TGCT), driven by aberrant CSF-1 production (Gelderblom, Bhadri, et al., 2024; Tap et al., 2019). Outside this context, monotherapy with edicotinib or LY3022855 has shown limited clinical benefit (Autio et al., 2020; von Tresckow et al., 2015). However, CSF1R inhibition causes consistent depletion of TAM in the TME, including pre-clinical models of CLL and follicular lymphoma (Galletti et al., 2016; Gomez-Roca et al., 2019; Valero et al., 2021). As emerging data highlights the strongly immuno-suppressive function of TAM on T cell function, CSF1/CSF1R inhibition mediated TAM depletion gains increasing interest to leverage efficiency of T cell immunotherapy. Pre-clinical models demonstrated that CSF1R inhibition augments CD8+T cell tumor infiltration and increased efficiency of immune-checkpoint inhibition in combination regimens (Peranzoni et al., 2018; Sato et al., 2025; Zhu et al., 2014). Moreover, Stahl et al. recently demonstrated that CSF1R inhibition effectively depletes TAMs and thereby strongly enhances CAR-T cell efficacy in a murine DLBCL model (Stahl et al., 2025). To date, trials combining T cell immunotherapy and CSF1R inhibition in solid tumors have yielded mixed outcomes (Foster et al., 2021; Razak et al., 2020; Weiss et al., 2021). However, further studies, including in hematological malignancies, are still needed. In CLL, a disease characterized by dependence on tumor supportive macrophages and low response to T cell immunotherapy resulting in resistance to checkpoint blockade, combining CSF1R inhibition with emerging immunotherapies such as CAR-T cells or bispecific antibodies might disrupt microenvironmental support and improve therapeutic outcomes (Lewis, Vom Stein, & Hallek, 2024). Our findings emphasize that alternative macrophage-depleting strategies, such as nanobody-based agents, merit further investigation as combination partners in future therapeutic approaches for CLL.

## Acknowledgements

This study was supported by the Deutsche Krebshilfe (DKH, German Cancer Aid) Foundation grant #70112403 to M.H. and by the Deutsche Forschungsgemeinschaft (DFG, German Research Foundation) grant SFB1530-455784452 (sub-project B01) to M.H. and P.H.N.. We thank Dr. E. Richard Stanley (NYC, US) for providing the *Csf1r* mouse and MacCsf1r cell lines and Dr. Emanuel Niesen for helpful discussion.

## Author contribution

Na.R. and G.R.R. analyzed data and wrote the manuscript. Na.R., V.K., T.T.T., A.F.v.S, M.K., S.R., performed experiments. S.I., Sh.I., and S.M. designed, synthesized and provided the Doxorubicin liposomes. N.R. designed the study and performed experiments. P.H.N. designed experiments, supervised the study and wrote the manuscript. M.H. initiated, designed and supervised the study. All authors revised and approved the manuscript.

