## Supplemental Data for "Reduced CSF1R expression in myeloid cells has limited impact on chronic lymphocytic leukemia progression"

### Supplemental Figure

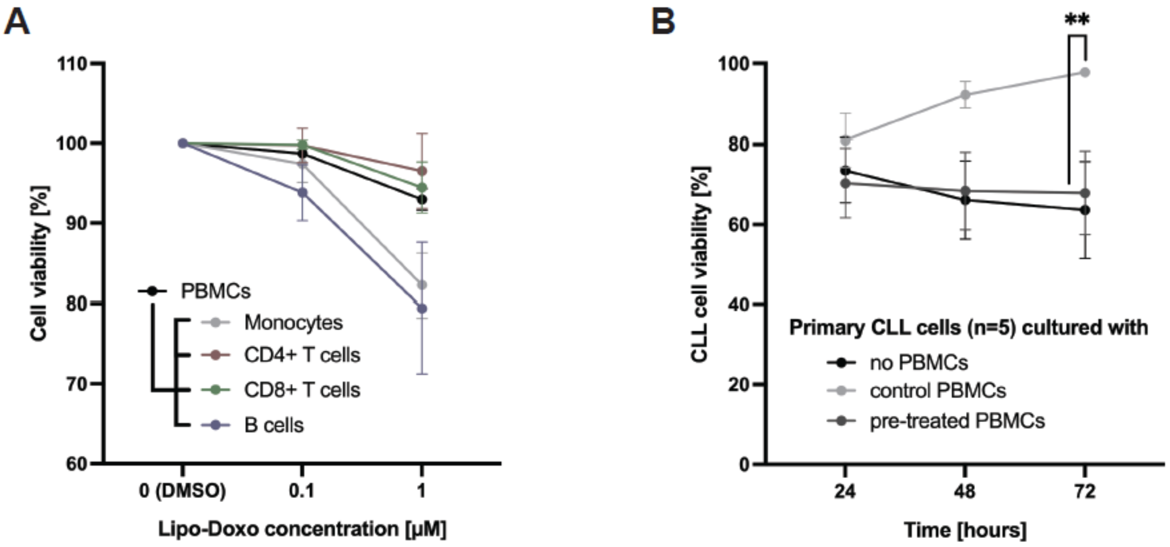


**Supplementary Figure 1. Validation of monocyte-dependent support of CLL cell survival.**
(A) Survival of total peripheral blood mononuclear cells (PBMCs) from healthy donors and distinct immune cell subsets within these PBMCs ($CD14^+$ monocytes, $CD4^+$ T cells, $CD8^+$ T cells, and $CD20^+$ B cells), after in vitro treatment for 24 hours with DMSO (control) or liposomal doxorubicin (Lipo-Doxo) at 0.1 µM or 1 µM. Cell viability was assessed by Annexin V staining and flow cytometry, and values were normalized to the respective DMSO control for each population. Statistical analysis was performed using the Wilcoxon test.
(B) Survival of patient-derived CLL cells co-cultured with healthy donor PBMCs that were pre-treated in vitro for 24 hours with DMSO or 1 µM Lipo-Doxo. Co-cultures were maintained for 24, 48 and 72 hours, after which the survival of CLL cells ($CD5^+$ $CD19^+$) was assessed as in (A). Non-normalized data were used for analysis.

### Supplemental Tables

**Table S1: List of fluorochrome labelled antibodies used for flow cytometry experiments.**

| Antibody | Manufacturer | Reference |
|---|---|---|
| CD5 | Miltenyi Biotec | Clone 53-7.3 |
| CD19 | Miltenyi Biotec | Clone 6D5 |
| CD45 | Miltenyi Biotec | Clone 30F11 |
| CD115 | Miltenyi Biotec | Clone AFS98 |
| CD115 Isotype control antibody (rat IgG2a) | Miltenyi Biotec | Clone ES26-15B7.3 |
| CD11b | BioLegend | Clone M1/70 |
| F4/80 | BioLegend | Clone BM8 |
| Ki-67 | BioLegend | Clone 16A8 |
| Ki-67 Isotype control antibody (Rat IgG2a) | BioLegend | Clone RTK2758 |

**Table S2: List of antibodies used for immunohistochemistry staining.**

| Antibody | Manufacturer | Reference |
|---|---|---|
| CD68 | Abcam | ab125212 |
| CD45R | BD Bioscience | Clone RA3-6B2 |

**Table S3: List of primary and secondary antibodies used in Western Blot analyses.**

| Antibody | Manufacturer | Reference |
|---|---|---|
| CD115 (CSF1R) | Cell Signaling | #3152 |
| GAPDH (D16H11) | Cell Signaling | #5174 |
| IRDye® 680LT anti-rabbit IgG Donkey | LI-COR | #926-68023 |
